# HSV-1 infection induces brain cofilin hyperphosphorylation in the 5×FAD Alzheimer’s Disease mouse model

**DOI:** 10.1101/2025.08.10.669568

**Authors:** Laia Gharavi-naeini, Zhenfeng Shu, Farhang Alem, Shu Feng, Linda D. Chilin, Pinghui Feng, Yuntao Wu

**Affiliations:** George Mason University, Center for Infectious Disease Research, School of Systems Biology, Manassas, VA; Section of Infection and Immunity, Herman Ostrow School of Dentistry, Norris Comprehensive Cancer Center, University of Southern California, Los Angeles, CA, USA

**Keywords:** Cofilin, Alzheimer’s disease, HSV-1, actin, LIM kinase, β-amyloid, cofilin-actin rods

## Abstract

Alzheimer’s disease (AD) is a degenerative neurological disease characterized by various biological signatures, including synaptic dysfunction, β-amyloid plaques, hyperphosphorylated Tau, cofilin-actin rods, and Hirano bodies, all of which are linked to the actin cytoskeleton and its regulators. Additionally, the presence of herpes simplex virus type 1 (HSV-1) in the brains of AD patients has long been suggested as a contributing factor for AD. However, mechanisms by which HSV-1 accelerates AD pathogenesis remain poorly understood. Here we report that HSV-1 infection induces hyperphosphorylation of cofilin in the brains of 5×FAD mice. Cofilin is an actin depolymerizing factor, and its S3 phosphorylation inactivates cofilin’s activity to depolymerize actin filaments. These findings facilitate the understanding of impacts of HSV-1 infection on the development of Alzheimer’s disease and have implications in AD therapeutics.

## Introduction

Alzheimer’s disease (AD), characterized by cognitive impairment, affects approximately 55 million people worldwide in 2020 (1), and currently there is no effective treatment. The accumulation of β-amyloid (Aβ) and hyperphosphorylated Tau in the brain are two major hallmarks of AD (2). Other molecular and cellular signatures of AD include the development of cofilin-actin rods (3) and Hirano bodies (4), all of which are related to the actin cytoskeleton and its regulators. Among actin-binding proteins, cofilin is the major actin depolymerizing factor that is ubiquitously expressed in the brain and excitatory synapses (5). Functionally, cofilin directly binds and depolymerizes filamentous actin (F-actin), and is responsible for the high turnover rates of actin filaments (6). The biological activity of cofilin is mainly regulated by phosphorylation of serine 3 at the N-terminus, which inhibits cofilin binding to actin. The kinases responsible for cofilin serine 3 phosphorylation are the LIM domain kinases (LIMK) and Tes kinases (7, 8). Cofilin is also activated through dephosphorylation of serine 3 by phosphatases such as PP1, PP2A, slingshot phosphatase 1 (SSH1), and chronophin (9-11).

Cofilin plays a critical role in the dynamics of dendritic spines, influencing their formation and elimination in response to synaptic activity (12-14). As such, cofilin dysregulation has been linked to synaptic dysfunction, which precedes AD pathogenesis (15). For instance, synaptic transmission is impaired with the loss of dendritic spines (16). This reduction in spines can directly affect cognitive functions, and is often observed in the postmortem brain tissues from AD patients and in primary neurons from AD mice (16-18). Cofilin-1 knockdown in neuronal cells led to decreases in both the number of thin spines and the length of dendritic protrusions (12).

Cofilin has also been suggested to be involved in cellular pathogenic responses to Aβ, and cofilin activity can be altered by the presence of β-amyloid aggregates. For example, it has been shown that Aβ_1-40_ and Aβ_25-35_ fibrils induce the activation of LIMK and cofilin S3 phosphorylation/inactivation (19). Paradoxically, other studies have also shown that cofilin can be activated by Aβ_1-40_ and Aβ_1-42_ in a LIMK-independent manner, which may be linked to the SSH1 pathway (20). Interestingly, it has been demonstrated that cofilin-1 phosphorylation /inactivation was increased with age and AD pathology, both *in vivo* and *in vitro*, and these changes were associated with inactivation of SSH1 (21).

Hyperphosphorylation of Tau, a microtubule-associated protein, is another hallmark of AD pathology. This hyperphosphorylation leads to the formation of neurofibrillary tangles and is associated with neuronal degeneration (22). Activated cofilin has been shown to interact with tubulin (23), and compete with Tau for direct microtubule binding both *in vitro* and *in vivo*, which inhibits Tau-induced microtubule assembly (24). In Tau-P301S mice, cofilin knockdown mitigates tauopathy and synaptic defects (24). Thus, cofilin could disrupt the normal function of Tau in stabilizing microtubules, leading to further destabilization of neuronal structure and function.

Cofilin’s abnormal interaction with actin can lead to the formation of pathogenic rod-shaped bundles of filaments, known as cofilin-actin rods (13), which occur in a 1:1 ratio of cofilin to actin in cultured primary neurons (25, 26). These rods have been observed in the brains of individuals with AD, particularly in the frontal cortex and hippocampus (3), as well as in transgenic AD mouse models (27). The presence of cofilin-actin rods is also considered as a pathological feature of AD, as treatment with Aβ_1-42_ can induce these rods in about 20% of cultured rat hippocampal neurons (27). Additionally, synthetic Aβ_1-42_ oligomers can gradually induce the formation of cofilin-actin rods in human cortical neurons (28).

Another similar structure, Hirano bodies, is frequently found in the Sommer’s sector of Ammon’s horn, where AD-associated Pick bodies and neurofibrillary tangles are also present (4). Hirano bodies primarily consist of ADF/cofilin and actin and are significantly elevated in AD patients (29, 30). The emergence of Hirano bodies in the Sommer’s sector may contribute to cognitive impairments observed in neurodegenerative disorders like AD.

Cofilin is also a common target for various viruses during infection. It has been proposed that the cortical actin serves as a natural barrier to viral entry and intracellular migration (31). Different viruses have evolved strategies to manipulate cortical actin for successful infection. For instance, in HIV-1 latent infection of blood resting T cells, the virus engages chemokine coreceptors CXCR4 and CCR5, activating G-protein signaling and cofilin to enhance the cortical actin dynamics needed for viral entry and nuclear migration (31-35). This dependency on cofilin and actin dynamics in viral infection is seen across multiple viruses, including herpes simplex virus (HSV-1) (36), alphaherpesvirus (37), vaccinia virus (38), measles virus (39, 40), Zika virus (41), and wheat dwarf virus (42). Notably, HSV-1 infection in neuronal cells results in biphasic cofilin and actin dynamics (36). In fact, the alphaherpesviral serine/threonine kinase US3 has been shown to promote significant cofilin dephosphorylation and restructuring of the actin cytoskeleton, aiding viral infection (37).

Potential contribution of HSV-1 infection to AD has long been speculated, as HSV-1 DNA has been detected in the brain of a significant number of AD patients (43), particularly in the temporal cortex and hippocampus regions (44), as well as within amyloid plaques (45). Population-based anti-HSV serological studies have further suggested that chronic HSV infection and viral reactivation are highly correlative with the incidence of AD (46). It has also been proposed that HSV-1 is a significant risk factor for AD in individuals who carry the type 4 allele of the apolipoprotein E gene (APOE-ε4) (47). Previously, it was proposed that HSV-1 glycoprotein B induces the nucleation of Aβ that acts as a defensive antimicrobial peptide against HSV-1 (48-50). However, a direct mechanistic linkage between HSV-1 infection and the development of the AD has yet to be firmly established. Given the role of cofilin in AD pathogenesis and the capacity of HSV-1 to modulate cofilin, we hypothesized that HSV-1 reactivation-induced dysregulation of cofilin in the brain could potentially connect HSV-1 and AD. In this study, we investigated HSV-1-mediated cofilin dysregulation in the brain using the 5×FAD AD mouse model.

## Results and Discussions

To recapitulate features of AD, we used female 5×FAD mice that carry human APP and PSEN1 transgenes (with a total of five-AD-linked mutations) (51). The 5×FAD mice present AD phenotypes relatively early. In particular, amyloid pathology can begin as young as 8 weeks of age and is more severe in females than in males (52). Mice were infected with HSV-1 at 12 weeks of age and were followed until 32 weeks old. The brain tissues were examined to determine the effects of HSV-1 on cofilin phosphorylation in infected and mock-infected control mice. Brain tissue sections were stained with a primary anti-phospho-cofilin (p-S3) antibody, followed by staining with a fluorescently-labeled secondary antibody for fluorescent microscopic imaging. Representative images from 3 independent experiments are shown in **Fig. 1**. We observed marked increases in the levels of p-cofilin staining in the brain of HSV-1-infected mice in comparison with mock-infected controls. Given that S3 phosphorylation leads to the inactivation of cofilin, this HSV-1-mediated increase in cofilin phosphorylation in the brain may impact AD pathogenesis in the genetic background of AD. These results may also suggest that HSV-1-induced cofilin hyperphosphorylation in the brain may serve as a possible molecular linkage connecting HSV-1 infection with AD.

**Figure 1.**
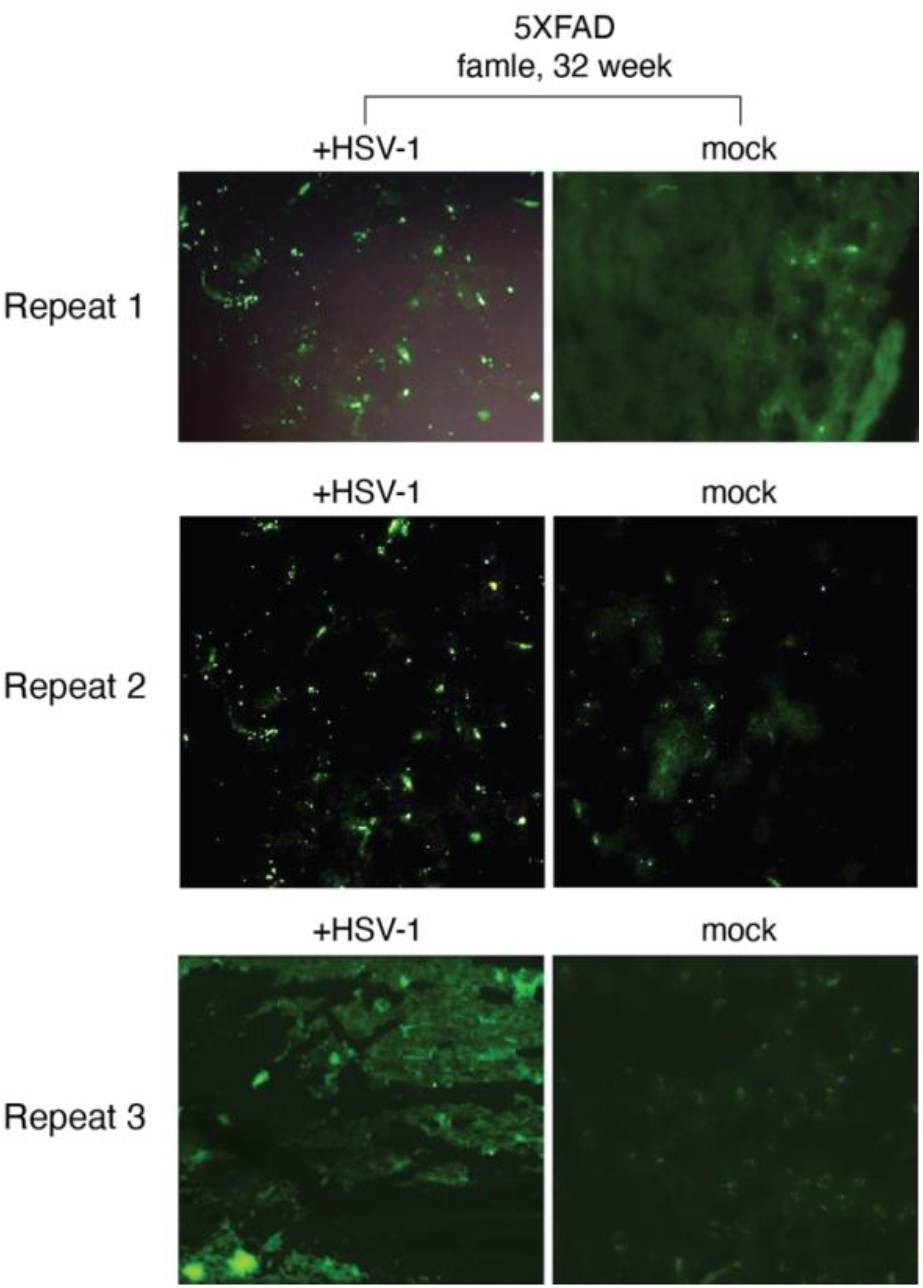
HSV-1 infection induces cofilin S3 hyperphosphorylation in the brains of 5×FAD mice. 5×FAD transgenic mice (12 week-old) were infected or mock-infected with HSV-1. Following infection, at 32 weeks of age, the brain tissues were examined to determine the effects of HSV-1 on cofilin phosphorylation by staining with a primary anti-phospho-cofilin (p-S3) antibody, followed by staining with a fluorescently-labeled secondary antibody for fluorescent microscopic imaging. Representative images from 3 independent experiments are shown.

We did not examine changes of total cofilin in HSV-1-infected mice, because previous studies have demonstrated that HSV-1 mainly alters cofilin phosphorylation rather than the total amount of cofilin in cells (36). HSV-1 triggers cofilin activity may be through viral kinases such as US3 (37). However, in cell culture conditions, US3 promotes cofilin activation rather than phosphorylation/inactivation with unknown mechanisms (37). An increase in cofilin phosphorylation can be achieved either by increasing the LIM kinase activity (33, 53) or by decreasing the activities of cofilin phosphatases such as PP1, PP2A, SSH1, and chronophin (20, 21). A detailed regulatory mechanism of HSV-1-induced cofilin hyperphosphorylation, as seen in our study, awaits to be determined.

In addition to HSV-1, another herpes virus, varicella zoster virus (VZV), has also been implicated in AD, and vaccination against shingles caused by VZV in senior citizens has been found to decrease the risk of AD/dementia (54-58). Interestingly, although VZV infection does not directly lead to Aβ and phospho-Tau accumulation in cells, VZV infection causes reactivation of HSV-1 and consequent AD-like changes, including Aβ plaque and phospho-Tau accumulation in cells (59). Our results may imply that targeting of HSV-1 reactivation or HSV-1-induced cofilin hyperphosphorylation in the brain may impact AD pathogenesis.

## Materials and Methods

All animal procedures were conducted in strict accordance with the recommendation in the Guide for the Care and Use of Laboratory Animals of the National Institutes of Health. The experimental protocol was approved by the Institutional Animal Care and Use Committee (IACUC) of the University of Southern California.

Age- and gender-matched 5×FAD transgenic mice (3-month-old) were used for mock and HSV-1 infection. Each mouse in the infection group was challenged with 5×10^5^ PFU of HSV-1 (strain 17) in 100 μl PBS via intravenous injection. Mock- and HSV-1-infected mice were sacrificed, and brains were collected for immunofluorescence staining, metabolite analysis, and gene expression analysis. Mouse frozen familial Alzheimer diseased (FAD) and wild type (WT) brain were stored in 4% PFA for more than 24 hours and transferred to 30% sucrose solution for another 24 hours. 10 μm thick brain sections on glass slides were prepared through cryostat sectioning. Slides were rinsed in 0.01 M PBS three times (5 min each time), followed by incubation with 0.5% Triton solution for 30 minutes to partially permeabilize the membrane and expose the antigen epitope, and then blocked with 5% bovine serum albumin containing 0.1% Triton for 1 hour. Next, diluted primary antibody, a rabbit anti-phospho-cofilin (Ser3) (77G2) monoclonal antibody (Cell Signaling) in 0.1% Triton X-100 in PBS (1:100 or 1:500) was added onto the sections according to manufacturer’s instructions, and incubated overnight at 4°C. On the next day, the sections were rinsed and incubated with diluted polyclonal anti-rabbit IgG (H+L) F(ab’)2 Fragment (Alexa Fluor-488-labelled) (Cell Signaling) (1:2000) for 2 hours at room temperature. Slides were rinsed and mounted in anti-fade mounting solution of 20% n-propyl gallate in DMSO. The fluorescence images were acquired by Nikon Ti2-E Widefield Fluorescence Microscope (Nikon Instruments Inc. Melville, NY) equipped with a 40x objective.

## Acknowledgments

The authors wish to thank the NIH AIDS Reagent Program for reagents. This work was supported by Public Health Service grant AI148012 (to Y.W.) and R56AI183995 (to Y.W., S.J., and D.K.K) and AG070904 (to P.F). Research in the Feng laboratory is supported by a generous startup fund from the Herman Ostrow School of Dentistry of USC.

